# Hyper-proliferation of Adipose Progenitors During Developmental Adipogenesis Programs Higher Adipocyte Number and Early-onset Obesity in Offspring Born to Obese Dams

**DOI:** 10.1101/2025.09.13.676029

**Authors:** TB Scheidl, JS Yoon, JL Wager, S Yonan, A Lakhdar, S Kailasam, JA Thompson

**Affiliations:** Cumming School of Medicine, University of Calgary, Calgary, AB; Department of Biochemistry and Molecular Biology, University of Calgary, Calgary, AB; Life Sciences and Biochemistry, Queens University, Kingston, ON; Canadian Core for Computational Genomics, McGill University, Montreal, QC

## Abstract

Being born to a mother who was obese during pregnancy is one of the strongest predictors of early onset obesity and metabolic syndrome. To identify the developmental mechanism linking maternal obesity to cardiometabolic disease in the offspring, a high fat/high fructose diet or low-fat control diet were introduced 4 weeks prior to mating in C57BL6 females and continued throughout pregnancy and lactation. Offspring born to obese dams had greater whole-body adiposity prior to puberty and were more susceptible to diet-induced obesity in adulthood. On postnatal day 10 (PND10) when pups born to obese dams had greater accumulation of subcutaneous adipose tissue, single cell sequencing was used to identify adipose progenitor cells (APCs) in the stromal vascular fraction. Two distinct lineages of APCs were identified that arose from a common proliferative root, terminating in committed preadipocytes and anti-adipogenic APCs. In pups born to obese dams, the proliferative root made up a greater proportion of APCs, more of these cells were in the G2M and S phases of the cell cycle and there was an enrichment in genes associated with proliferation (*mKi67, Fat3, Ccn4, Gata4*). On PND10, APCs isolated from the inguinal depots of pups born to obese dams were hyper-proliferative and there was a greater abundance of total APCs, whereas there were no differences in adipocyte size. After puberty, APCs in subcutaneous depots of offspring born to obese dams were more adipogenic in vitro and this group gained a greater amount of body fat in response to pharmacological stimulation of adipogenesis by rosiglitazone. Male adults born to obese dams had a lower abundance of APCs and were more susceptible to APC exhaustion due to prolonged exposure to rosiglitazone. Therefore, early onset obesity in offspring exposed to maternal obesity in utero is due to a prolongation of the proliferative phase of early life adipogenesis, leading to a higher adipocyte number.

## INTRODUCTION

Obesity rates among reproductive aged women have climbed over past decades, with approximately 40% of women now estimated to be classified as obese [1]. In the setting of pregnancy, the impact of obesity is not restricted to the pregnant mother but is transmitted to the fetus during sensitive windows of development. Being born to a mother who is obese during pregnancy increases the odds of childhood obesity by 3-fold [2–4] and children with high adiposity or obesity during childhood are 77% more likely to remain obese as adults [5]. Evidence in both human populations and animal studies demonstrate that this familial cycle of obesity is in part attributable to the intrauterine environment independent of genetic susceptibility [6–8]. Dissecting these environmental causes from the genetic component of trans-generational cycles of obesity is critical as the former can be targeted for primary prevention.

The primary outcome of pregnancies complicated by obesity is fetal macrosomia, which is diagnosed when birth weight is large-for-gestational age and characterized by elevated adiposity. Thus, a perturbation in the development of white adipose tissue (WAT) is likely an initiating event underlying obesity predisposition in the offspring [9]. Developmental establishment of subcutaneous adipose depots, which constitute ∼ 90% of total white adipose tissue (WAT) volume, starts in fetal life, whereas development of the secondary, visceral depots is largely a postnatal event. At E14.5, mesenchymal stem cells are directed into the adipocyte lineage and undergo a period of rapid proliferation followed by terminal differentiation into adipocytes that coincides with a peak in fat accretion during the first week of life. Following a second phase of adipose progenitor cell (APC) proliferation and fat accumulation in puberty, adipocyte numbers remain stable throughout life. Post-pubertal consistency in adipocyte number is owing to a balanced cycle of adipocyte death and birth, and was demonstrated across a range of BMIs in human adults by Spalding *et al* [10]. This landmark study revealed a higher adipocyte number in adults with early-onset obesity, suggesting that adipocyte number is established early in life and is predictive of childhood obesity [10]. Therefore, early life adipogenesis establishes the set-point of adiposity and may underlie the developmental origin of early onset obesity in offspring exposed to maternal obesity *in utero*.

In adulthood, subcutaneous WAT depots serve as the body’s metabolic sink and play a central role in regulating whole-body metabolic homeostasis. The expansion of WAT to accommodate increased demands for energy storage preserves metabolic health and distinguishes metabolically healthy from metabolically unhealthy obesity. While WAT expansion in adulthood relies primarily on adipocyte hypertrophy, new lines of evidence implicate a resident population of APCs as critical for maintaining the lipid buffering capacity of WAT under obesogenic conditions. In rodent models of obesity [11], impaired adipogenesis is associated with worse metabolic health, while stimulating adipogenesis preserves metabolic health [12]. Intriguingly, Jiang *et al*. elegantly demonstrated using a PPARᵧ-dependent fate mapping model that APCs residing in adult depots are a distinct lineage that undergo fate determination earlier in gestation compared to the APCs that give rise to adipocytes during early life adipose development [13]. Therefore, intrauterine life is a critical window that drives adipose organogenesis and influences the availability and phenotype of APCs in adulthood. Thus, a suboptimal intrauterine environment may alter determination of progenitors to the adipose lineage and thereby influence life-long adiposity and the ability of WAT to respond to fluctuations in energy balance.

Herein, we sought to determine if heightened vulnerability to metabolic syndrome in children born to obese mothers begins with a perturbation in the developmental establishment of adipose tissue.

## MATERIALS AND METHODS

### Animals

All experiments involving animals were performed at the University of Calgary with approval from the institutional animal care committee and conducted in accordance with guidelines from the Canadian Council on Animal Care Ethics. Virgin female C57BL/6J mice were fed a low-fat diet (LFD, 10% kCal from fat) or high fat/high fructose diet (HFFD, 35% kCal from fructose, 45% kCal from fat) (table 1) for 4 weeks prior to pairing with experienced studs and throughout pregnancy and lactation. Animals were anesthetized by inhalation of 5% isoflurane followed by humane euthanasia by decapitation. Inguinal WAT (iWAT) or gonadal WAT (gWAT) was collected for subsequent experiments. Whole blood was obtained at euthanasia for serum collection.

**Table 1.** Experimental Diet Composition.

| Diet Name | Macronutrient Composition<br>(%gm) |  |  | Micronutrient<br>Composition (%gm) |  |
| --- | --- | --- | --- | --- | --- |
|  | Fat | Carbohydrate | Protein | Vitamin | Mineral |
| **High Fat/High Fructose Diet (45% kCal from fat, 35% kCal from fructose)** | 23.6% | 41.4% | 23.7% | 10% | 10% |
| **Low Fat Diet (10% kCal from fat)** | 4.3% | 66.7% | 21.8% | 0.3% | 4.7% |

#### Postnatal Interventions

To induce obesity, adult offspring of lean or obese dams were fed LFD or HFFD from the beginning of adulthood (7 weeks of age) for a total of 16 weeks, as demonstrated previously [8, 14]. In a separate subset of offspring, adipogenesis was pharmacologically induced, beginning at 12 weeks of age, by rosiglitazone (rosi, 15mg/kg/day) dissolved in 1% methylcellulose and diluted in 50% sweetened condensed milk daily for 8 weeks. Controls were administered 1% methylcellulose in 50% sweetened condensed milk as a vehicle.

### Assessment of Body Composition and Glucose Tolerance

Whole body fat and lean mass were measured by time domain nuclear magnetic resonance (TD-NMR). Crown to rump length was measured using electronic calipers. Blood glucose was measured in 12-hour fasted animals using a commercially available glucometer.

### Isolation of Primary APCs

Under sterile conditions, iWAT was collected for isolation of the stromal vascular fraction (SVF) as previously described [8]. The resultant SVF pellet containing APCs was used for subsequent experiments.

### Single Cell RNA Sequencing (scRNA-seq)

For scRNA-seq profiling of APCs, timed pregnancies for lean and obese dams were carried out such that dams from each group delivered on the same day. The SVF was collected from two PND10 offspring per litter, which was subsequently repeated. Isolated SVF was filtered to generate a single cell suspension. Cell viability was determined prior to library preparation using the Invitrogen Countess 3 FL Automated Cell Counter and the ReadyCount Green/Red Viability Stain. Single cell libraries were prepared by the University of Calgary’s Center for Health Genomics and Informatics using the 10x Genomics Chromium Next GEM Kit v.3.1 (PN: 1000158) according to the manufacturer’s instructions. Libraries underwent paired-end sequencing on the Illumina NovaSeq 6000 with 1% PhiX using the following cycle parameters: 28 (read 1), 90 (read 2), 10 (index i7), 10 (index i5).

### Bioinformatics Analysis

Sequencing outputs were converted into FASTQ files, demultiplexed, and barcodes were counted with the 10x Genomics Cell Ranger software v.8.0.1. Downstream analyses were performed using the R package Seurat v.5.3.0 [15]. Only cells that expressed 300-8000 genes (nFeature_RNA), 500-50000 transcripts (nCount_RNA), and less than 30 percent mitochondrial genes were retained in the dataset. Seurat objects were created for each sample, which had any doublets removed with DoubletFinder [16] and integrated with Harmony [17]. Clustering, dimensional reduction, differentially expressed gene (DEG) analysis, and visualizations were performed in Seurat. Cutoffs for all statistically significant DEGs were adjusted p-value < 0.05 using the Wilcoxon rank-sum test. Pseudotime and trajectory analysis were performed in Monocle3 with cell embeddings from Seurat [18–21]. Gene ontology analysis was performed with enrichR v.3.4 [22–24].

### APC Culture and Differentiation

At first passage, APCs were cultured for assessment of *in vitro* differentiation by lipid droplet staining and gene expression. Seeded cells were monitored until reaching 100% confluency and held at confluency for 48h to reach contact inhibition (D0) [25]. Cells were grown and differentiated using commercially available adipocyte differentiation media.

### Measurement of *in vitro* Adipogenesis and Lipid Droplet Accumulation

Cultured APCs were stained with Oil Red O for identification of lipid droplets after 4 (D4) or 7 (D7) days of differentiation, as previously described [8]. A subset of fixed cells was stained with 2mM BODIPY and 10mM Hoescht 33342 for 30 minutes to acquire representative images using the 10X objective on a Nikon TS2 brightfield microscope.

### Quantification of *in vitro* Proliferation

Upon passage, cells were seeded at 20,000 cells/mL. Cells were allowed to adhere for 4 hours and then pulsed with 10mg/mL BrdU overnight, after which cells were stained for flow cytometric quantification of BrdU using the eBioscience BrdU Staining Kit for Flow Cytometry. BrdU incorporation was quantified using non-BrdU pulsed cells as a gating control. Samples were analyzed using the Cytek Aurora Spectral Flow Cytometer and FlowJo 10.9.0 analysis software as described above.

### Determination of APC Abundance by Flow Cytometry

Isolated SVF was resuspended at 0.5×10^6^ cells/mL and stained with fixable viability dye concurrently with fluorescently tagged antibodies for 1 hour on ice (table 2). Following antibody staining, samples were fixed in 1% PFA for 15 minutes and stored at 4°C until analysis. Samples were run on the Cytek Aurora Spectral Flow Cytometer and single cells identified based on forward- and side-scatter (FSC, SSC). Endothelial cells and leukocytes were excluded by CD31 and CD45, respectively. Live CD31^-^/CD45^-^ (Lin^-^) cells that were doubly positive for CD34 and PDGFRα were identified as APCs. Fluorescent minus one (FMO) controls were used to distinguish between positive and negative events. Data were analysed using FlowJo version 10.9.0 and are represented as percentage of live cells.

**Table 2.** Antibodies for Flow Cytometric Quantification of APCs.

| Antibody Target | Fluorochrome | Volume for<br>0.5x10<sup>6</sup> Cells | Manufacturer |
| --- | --- | --- | --- |
| **CD45** | FITC | 2μL | Invitrogen (11-0451-85) |
| **CD31** | PE-Cy7 | 5μL | Invitrogen (25-0311-82) |
| **CD34** | Alexa Fluor 647 | 4μL | Biolegend (119314) |
| **Pdgfrα** | PE | 2.5μL | Biolegend (135906) |
| **CD24** | Percp Cy5.5 | 1.34μL | BD Pharmigen<br>(562360) |

### RNA Isolation and Measurement of Gene Expression

RNA was isolated from undifferentiated (D0) and differentiated (D7) APCs isolated from offspring of lean or obese dams. RNA was isolated using the Qiagen RNeasy kit according to manufacturer instructions, quantified by nanophotometer, and assessed for integrity using TapeStation assay (University of Calgary Genomics). Complementary DNA (cDNA) was synthesized from 1.6µg of RNA using the Applied Biosystems High-Capacity cDNA reverse transcription kit. Gene expression was measured by RT-qPCR with the Powerup SYBR Green Master Mix and the QuantStudio 5 Thermocycler under the following conditions: 50°C x 2 min, 95°C x 2 min, 95°C x 1 sec, 58.9°C x 30s (40 cycles).

Primers were designed using NCBI primer blast. Amplification was performed in triplicate and fold change calculated relative to *Actb* (β-actin) expression.

### Histological Analysis of WAT

Fixed WAT was embedded, sectioned, and stained for hematoxylin and eosin (H&E) by the University of Calgary Diagnostic Services Unit. Stained tissues were imaged using the Nikon TS2 brightfield microscope or the Olympus VS120 slide scanning microscope. The maximum diameter of each adipocyte was determined using QuPath (version 0.6.0). The proportion of adipocytes in at discrete size ranges (>10um) in each image was calculated.

### WAT Whole Mount Clearing, Staining and Imaging

Fixed ∼3mm^3^ sections of iWAT from PND10 offspring were stained with Lipidtox Deep Red (ThermoFisher), for 3 days at 4°C for representative images of adipocytes. Stained samples were placed on coverslip bottom 35mm dishes with a small amount of fluorescence mounting medium and imaged using the Leica SP8 confocal microscope with the 10X objective.

### Circulating Markers of Metabolic Health and Adipokines

Luminex® xMAP® technology was used to quantitatively detect twelve murine metabolic hormones and peptides. The multiplexing analysis was performed by Eve Technologies Corporation (Calgary, Alberta, Canada) using the Luminex® 200™ system (Luminex Corporation/DiaSorin, Saluggia, Italy) with Bio-Plex Manager™ software (Bio-Rad Laboratories Inc., Hercules, California, USA). Twelve markers were measured in the samples using the Eve Technologies’ Mouse/Rat Metabolic Hormone Discovery Assay® Array (MRDMET12) as per the manufacturer’s instructions (MILLIPLEX® Mouse Metabolic Hormone Expanded Panel Cat. # MMHE-44K, MilliporeSigma, Burlington, Massachusetts, USA). The 12-plex consisted of Active Amylin, C-Peptide, Active Ghrelin, Total GIP, Active GLP-1, Glucagon, Insulin, Leptin, PP, PYY, Resistin and Secretin. Assay sensitivities of these markers range from 1.4 – 91.8 pg/mL. Individual analyte sensitivity values are available in the MilliporeSigma MILLIPLEX® protocol. Data are shown only for insulin, leptin, and resistin. Free fatty acids were quantified per manufacturer’s instructions using the Abcam free fatty acid kit (ab65341).

### Quantification of Adipose Tissue Macrophages

Macrophages were quantified in the SVF of digested WAT following a slightly modified isolation protocol from above. Briefly, following digestion, the floating adipocyte layer was removed and cells pelleted. Cells were resuspended and filtered through a 100μm tissue strainer. One million cells/mL were stained with viability dye and fixed in 2% PFA, followed by antibody staining (table 3). M1 macrophages were classified as CD11c^+^/CD206^-^, and M2 macrophages as CD11c^-^/CD206^+^. After staining, samples were washed and analyzed on the Cytek Aurora Spectral Flow Cytometer as described above.

**Table 3.** Antibodies for Flow Cytometric Quantification of WAT Macrophages.

| **Antibody Target** | **Fluorochrome** | **Volume for**<br>**1X10<sup>6</sup> Cells** | **Manufacturer** |
| --- | --- | --- | --- |
| **CD45** | BV510 | 5µL | Biolegend (103151) |
| **F4/80** | APC/Fire 810 | 5µL | Biolegend (123166) |
| **CD11B** | FITC | 2µL | Biolegend (101206) |
| **CD11C** | PerCp | 3.5µL | Biolegend (117326) |
| **CD206** | Alexa 700 | 2µL | Biolegend (141734) |

### Statistical Analyses

Single variable comparisons between two groups were made using a students unpaired t-test. Single variable comparisons of three or more groups were performed with one-way ANOVA followed by Fisher’s multiple comparisons test. Multiple variable analyses were made using two-way ANOVA with Tukey’s multiple comparisons post-hoc test. Comparisons were considered significant when p<0.05. For transcriptomic comparisons in scRNA-seq data, comparisons were considered significant when there was a fold change ≥2 and p<0.05.

## RESULTS

### *In Utero* Exposure to Maternal Obesity Programs High Adiposity in Early Life and Throughout the Lifespan

Obese dams had significantly greater whole-body fat mass prior to pregnancy and at the end of weaning, compared to lean dams (Supp. Fig. 1A, B), with no difference in gestational weight gain (Supp. Fig. 1C). Obese dams had elevated 12-hour fasting glucose and a blunted post-prandial glucose response (Supp. Fig 1D – F). Litter size was not different between lean and obese pregnancies (Supp. Fig. 1G). Offspring adiposity and body weight were measured in the early postnatal period (PND 1 – 10), throughout puberty (PND 14 – 35) and during LFD or HFFD feeding in adulthood (>7 weeks) (Fig. 1A). Postnatal body weight was similar between the two offspring groups until PND20, at which point body weight gain plateaued in offspring of lean dams until weaning (PND21), whereas in offspring of obese dams, body weight continued to climb steadily until weaning (Fig. 1B). At weaning (PND21), both male and female offspring of obese dams had significantly greater whole-body fat mass, iWAT mass, and crown-to-rump length (Fig. 1C – F), an index of developmental growth. The control group demonstrated the expected pubertal spurt in fat accumulation from PND21 – 35, while the percent change in body fat mass over the course of puberty was close to zero in male and female offspring of obese dams, which both started puberty at higher levels of adiposity (Fig. 1G – I, K). In adulthood, male and female offspring of obese dams were more susceptible to diet-induced obesity (Fig. 1L – N, Q – S). Even when on the LFD, female offspring of obese dams had higher whole-body adiposity vs. females born to lean dams (Fig. 1Q). In all offspring, expansion of both iWAT and vWAT contributed to diet-induced increases in whole-body adiposity (Fig. 1L – U). More severe obesity in males born to obese dams was driven by greater expansion of iWAT (Fig. 1O, P); whereas more severe obesity in females born to obese dams was driven by greater expansion in both depots (Fig. 1T, U). Therefore, our mouse model of maternal obesity recapitulates the developmental programming of early onset obesity reported in human studies. Early onset obesity was apparent prior to puberty and attributable to greater fat accretion over the first weeks of postnatal life, followed by a virtually absent pubertal surge in fat mass. Consistent with human data showing a tendency for childhood obesity to track into adulthood, offspring born to obese dams were more susceptible to diet-induced obesity once sexually mature.

**Figure 1.**
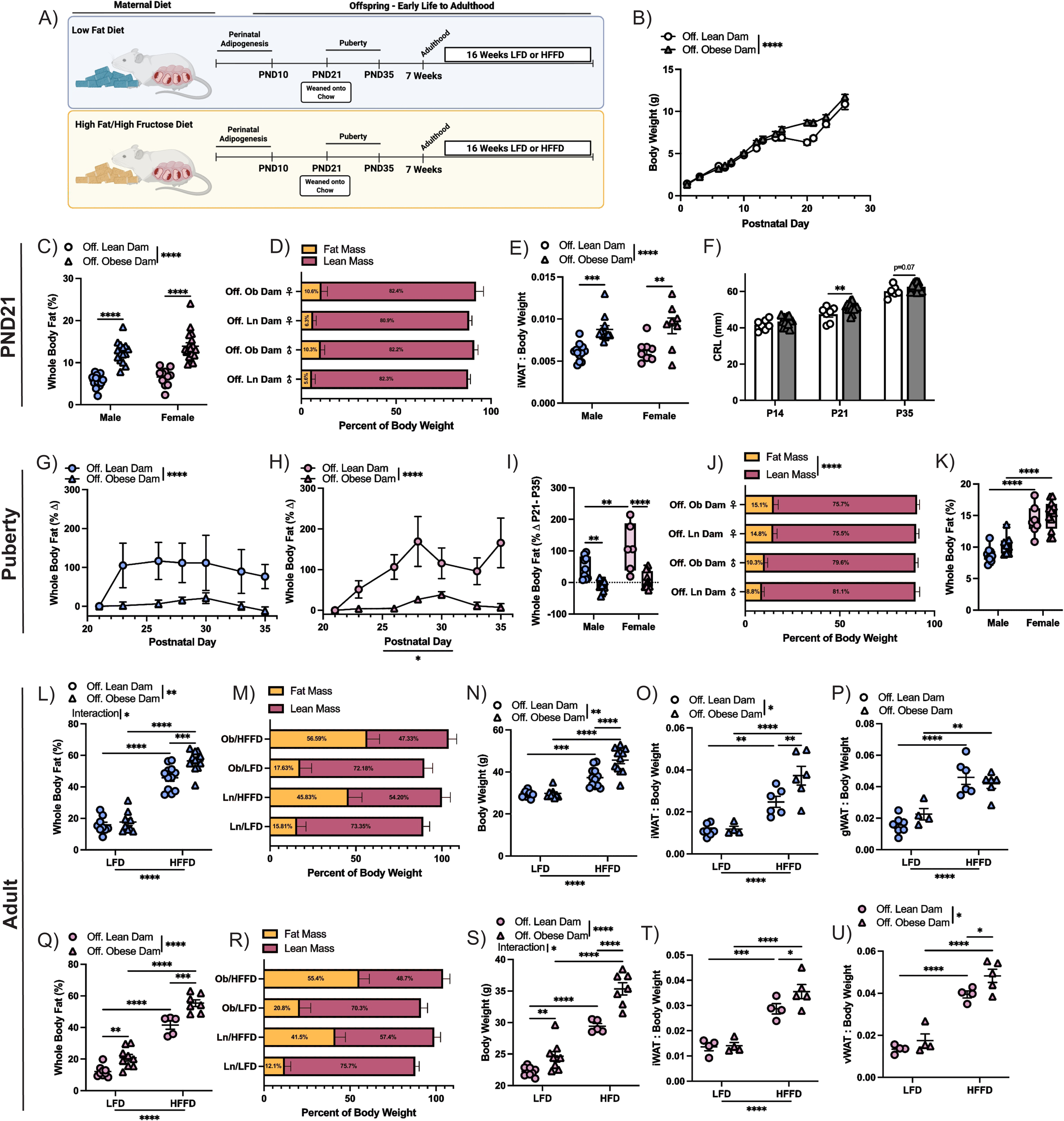
Developmental Exposure to Maternal Obesity Increases the Trajectory of Fat Accumulation Throughout the Lifespan. A) Schematic of experimental paradigm. Dams were fed LFD or HFFD prior to pregnancy, during pregnancy, and throughout lactation. Offspring were studied at PND10, 35, and after 16-weeks LFD or HFFD feeding beginning at ∼7 weeks of age. B) Body weight in offspring of lean vs. obese dams from PND1 – 26. Data for male and female animals are combined. C) Whole-body adiposity, determined by TD-NMR on PND21. D) Relative lean and fat mass in offspring on PND21. E) iWAT weight relative to body weight in offspring on PND21. F) Crown-to-rump length in offspring on PND14, 21, and 35. Data are combined for both sexes. Percent change in whole-body fat throughout puberty in (G) male and (H) female (H) offspring. I) Overall percent change in whole-body fat between PND21 and 35. J) Relative lean and fat mass in offspring at the end of puberty. K) Absolute whole-body fat mass after puberty. L) Percent whole-body fat mass and (M) relative percent fat and lean mass in male offspring at the end of LFD or HFFD. N) Body weight in male offspring at the end of LFD or HFFD feeding. O) Relative weight of iWAT and (P) gWAT depots in male offspring at the end of LFD or HFFD feeding. Q) Percent whole-body fat mass and (R) relative percent fat and lean mass in female offspring at the end of LFD or HFFD feeding. S) Body weight in female offspring at the end of LFD or HFFD feeding. T) Relative weight of iWAT and (U) gWAT depots at the end of LFD or HFFD feeding. *p<0.05, **p<0.01, ***p<0.001, ****p<0.0001.

### Single Cell RNA Sequencing Reveals an Actively Proliferating APC Population in SVF of PND10 Offspring

Given that adipocyte number is likely programmed during a developmental window of APC proliferation and differentiation spanning from late gestation into the first days of postnatal life, we profiled APCs in the SVF of iWAT at PND10 using scRNA-seq. SVF from two PND10 males was pooled into a single sample for library generation, for a total of four pups born to two pregnancies represented as two samples in each group (Fig. 2A). Following standard filtering and quality control steps, we obtained an integrated dataset of 20961 cells, composed of 12454 cells (59.4%) from the offspring of lean dams and 8507 cells (40.6%) from offspring of obese dams. Following initial clustering, clusters were annotated with their biological cell type based on canonical marker gene panels, which identified 10 expected cell types (Fig. 2B, C), with no significant differences in the proportion of cells from each group within a cell type (Supp. Fig. 2A). As expected, APCs (*Pdgfra, Cd34*, *Dlk1*, and *Ly6a)* comprised the largest proportion (40.1%) of the dataset with 8414 cells (Fig. 2D) and were subsetted for re-clustering and in-depth analysis.

**Figure 2.**
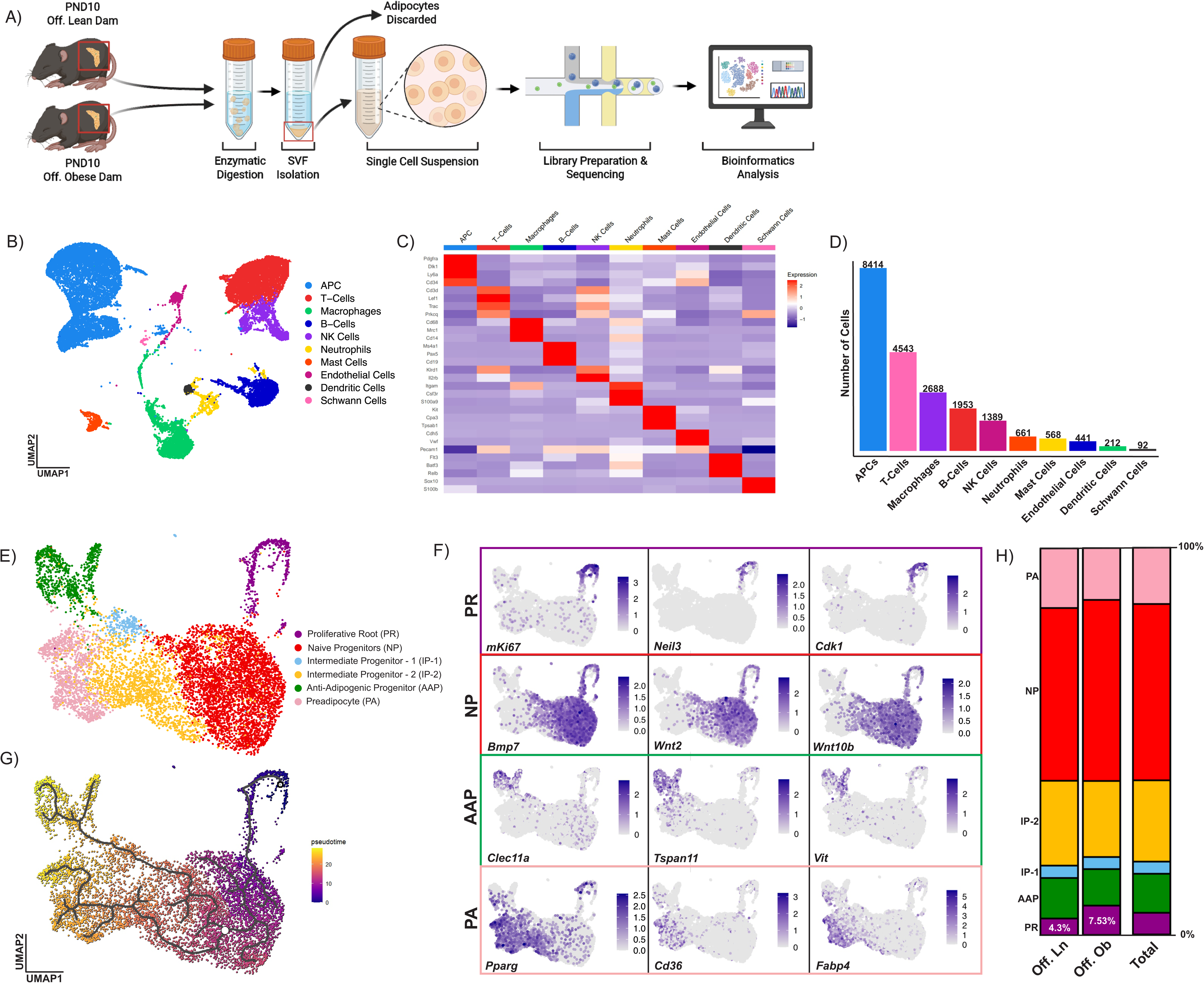
A Single-Cell Interrogation of the Composition of Stromal Vascular Fraction at PND10. A) Experimental design. Stromal vascular fraction was isolated from male offspring of lean or obese dams on PND10. Cells were evaluated for viability and then underwent 10X chromium library preparation and sequencing. B) Canonical cell clusters in SVF and C) marker genes for each cluster. D) Number of cells in each cluster. E) Curated APC clusters. F) Feature plots of marker genes for Proliferative Root, Naïve Progenitor, Anti-Adipogenic Progenitor, and Preadipocyte APC clusters. G) Pseudotime analysis of APC cluster. H) APC cluster proportion in offspring of lean vs. obese dams.

Further clustering identified 8 clusters of APCs (Supp. Fig. 2B) and unbiased DEG analysis was performed comparing each APC cluster against all other APC clusters. Clusters that did not express a distinct set of DEGs were merged for subsequent analysis, which resulted in 6 curated clusters (Fig. 2E). Interestingly, one cluster that we named the “Proliferative Root” (PR) uniquely expressed genes related to active proliferation, cell cycle, and DNA replication, such as *mki67, Cdk1,* and *Neil3* (Fig. 2F). We also found that 45.5% of APCs as a single large cluster were enriched for common mesenchymal stem cell (MSC) markers, including *Bmp7, Wnt2*, and *Wnt10b*, and therefore named this cluster “Naïve Progenitors” (NP) (Fig. 2F). A “Preadipocyte” (PA) cluster was identified by expression of common early adipocyte markers (*Pparg, Cd36,* and *Fabp4*) and an “Anti-Adipogenic Progenitor” (AAP) cluster by enrichment of alternative differentiation markers (*Clec11a, Tspan11,* and *Vit)* (Fig. 2F) [26]. Trajectory analysis identified two distinct lineages, with the proliferative root cells defined as the root node due to their high proliferation and MSC marker gene expression (Fig. 2G). The branch point of the two lineages occurred in the Naïve Progenitors, with one lineage terminating with Preadipocytes and the other with Anti-Adipogenic Progenitors. Two clusters that were named “Intermediate Progenitor – 1” (IP-1) and “Intermediate Progenitor – 2” (IP-2) bridged the mesenchymal stem cell root to the terminal preadipocyte and anti-adipogenic clusters (Fig. 2E) These Intermediate Progenitor clusters expressed markers of the Naïve Progenitors (*Dpp4, Wnt2, Pi16, Bmp7*), Anti-Adipogenic Progenitors (*F3/*CD142, *Clec11a*), and committed Preadipocytes (*Pparg, Fabp4*). *Cilp* exclusively marked the Intermediate Progenitor – 1 cluster (Supp. Fig. 2C).

### Interrogating Adipose Progenitor Cell Heterogeneity at Single Cell Resolution Reveals Naïve APCs from Offspring of Obese Dams are Hyperproliferative

While there were no differences between groups in the proportion of cells in the two lineages overall (Supp. Fig. 3A), there was a higher proportion of total APCs derived from offspring of obese dams that comprised the Proliferative Root cluster (54%) compared to offspring from lean dams (46%) (Fig. 2H, Fig. 3A). Thus, we compared DEGs between offspring of lean vs. obese dams within this cluster. Only 12 genes were significantly enriched in cells derived from offspring of lean dams, while 43 genes were enriched in cells from offspring of obese dams (Supp. Table 1). Of the 43 genes upregulated in cells from offspring of obese dams, the most significant DEGs with the greatest fold difference in expression were associated with cell cycle, proliferation, and DNA replication (*Pgrmc1, Pbm3, Aff3, Impdh2, Smoc2, Gab2, Wfdc1),* but were restricted to a relatively small subset of cells in the Proliferative Root cluster (Fig. 3B). To interrogate whether cells in the proliferative root cluster were indeed replicating, cell cycle scoring was performed. Unlike the majority of cells in other APC clusters, the majority of cells in the Proliferative Root cluster were in the G2M (62%) and S phase (33%) of the cell cycle (Fig. 3C, D). Additionally, *mki67*, a robust marker of proliferation, was almost exclusively expressed in the Proliferative Root cluster, though present within cells from both groups (Supp. Fig. 3C). To determine whether the Proliferative Root was a unique APC subtype or cells from another cluster that had entered the cell cycle, we compared DEGs in the proliferative root cluster vs. all other APC subclusters. Notably, there was almost complete overlap of marker genes identifying Naïve Progenitor cluster within the Proliferative Root cluster (Fig. 3E), but cells in the Naïve Progenitor cluster did not express genes distinguishing the Proliferative Root cluster (Fig. 3E). These data suggest that cells in the Proliferative Root cluster are Naïve Progenitors that remain in the cell cycle. Several genes (*mKi67, Fat3, Ccn4, Gata4*) that are involved in proliferation were almost exclusively expressed in Proliferative Root cells from offspring of obese dams (<1% expression in offspring of lean dams) (Supp. Fig. 3C), specifically in cells that were in the G2M and S phase of the cell cycle (Fig. 3F). Further, when comparing cell cycle stage composition in the proliferative root cluster between the groups, cells derived from offspring of obese dams had more cells in the G2M (55%) and S phase (54%) than cells from offspring of lean dams, suggesting that they were undergoing mitosis (Fig. 3G). Together, these data demonstrate that Proliferative Root cells are proliferating Naïve Progenitors, and that a greater number of these proliferating APCs exist in the iWAT of pups born to obese dams during the early postnatal period of adipogenesis.

**Figure 3.**
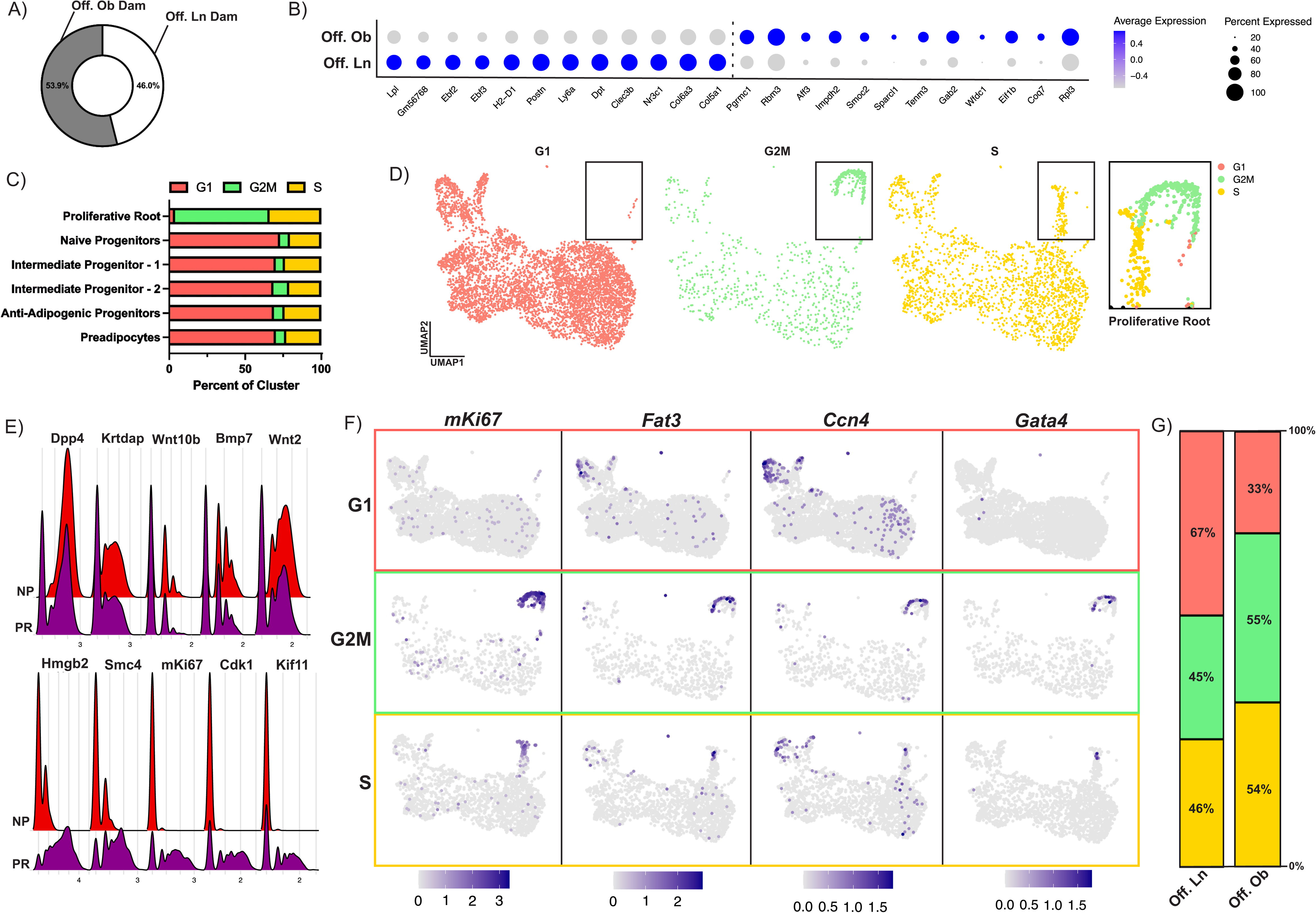
Proliferative Root APCs are Naïve APCs in Active Phase of Cell Cycle. A) Percent of cells in proliferative root contributed by offspring of lean or obese dams. B) DEGs in Proliferative Root cluster between offspring groups. C) Percent of cells scored in G1, G2M, and S phase within each APC cluster. D) Feature plots showing cell cycle score across APC cluster. Comparison of expression of marker genes in Proliferative Root and Naïve Progenitor cluster. F) Feature plots showing expression of genes related to proliferation in each cell cycle phase. G) Proportion of cells in each cell cycle phase within the Proliferative Root, compared between offspring groups.

### Accelerated Fat Mass Accumulation in Early Life Results from Increased APC Proliferation and Adipocyte Number

Given our transcriptomic evidence that APCs in PND10 offspring born to obese dams are more proliferative in early postnatal life, we next sought to determine if higher early life adiposity was attributable to generation of more adipocytes or greater lipid accumulation in differentiated adipocytes. On PND10, APCs isolated from iWAT of pups born to obese vs. lean dams were more proliferative in culture (Fig. 4A, B) and present in greater abundance in the SVF when identified by cell surface markers (CD34^+^/PDGFRα^+^) in flow cytometry (Fig. 4C, D). In line with a more proliferative APC phenotype at PND10, fewer APCs isolated from offspring of obese dams had differentiated into adipocytes during the early stage of in vitro differentiation (Fig. 4E), while during the maintenance phase there were no differences in lipid droplet accumulation between the two groups (Fig. 4F). The markedly higher subcutaneous adiposity at PND10 in pups born to obese dams (Fig. 4G, H) was not attributable to differences in adipocyte size (Fig. 4J) and there were no differences in serum leptin (Fig. 4K). These data demonstrate that the elevated setpoint of early life adiposity in offspring exposed to maternal obesity *in utero* is owing to APC hyper-proliferation and higher adipocyte number rather than a perturbation in lipid synthesis pathways.

**Figure 4.**
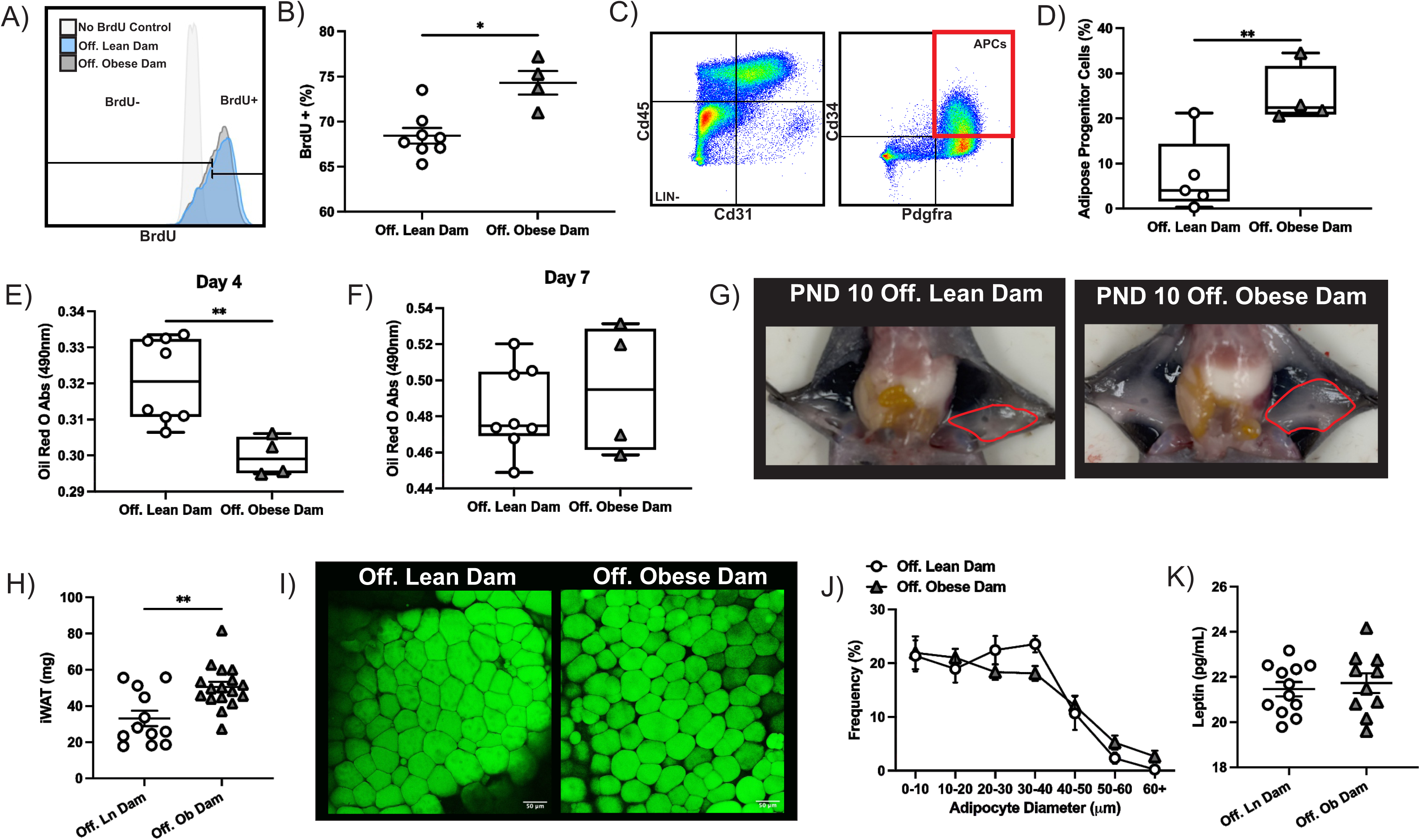
Subcutaneous APCs in Offspring of Obese Dams are Hyperproliferative. A) Gating paradigm for cultured APCs pulsed with BrdU. B) Flow cytometric quantification of BrdU^+^ APCs derived from PND10 offspring of lean vs. obese dams. C) Gating paradigm for quantification of APCs from iWAT of PND10 offspring. D) Abundance of CD34^+^/PDGFRα^+^ APCs in iWAT of PND10 offspring of lean vs. obese dams. Quantification of neutral lipid staining by Oil Red O in cultured APCs derived from PND10 offspring of lean vs. obese dams and after (E) 4 and (F) 7 days of differentiation. G) Representative image of iWAT depot in PND10 offspring of lean vs. obese dams. H) iWAT mass in PND10 offspring. I) Representative Lipidtox stained image of whole mount iWAT at PND10. J) Quantification of adipocyte size in iWAT in PND10 offspring. K) Serum leptin concentration in PND10 offspring. *p<0.05, **p<0.01.

### *In utero* Exposure to Maternal Obesity Predisposes to APC Exhaustion in Adult Offspring

Since APCs residing in established adult depots are distinct from developmental APCs in both lineage trajectory and function, yet also originate in utero, we set out to determine how exposure to maternal obesity impacts adult APC phenotype. Recruitment of adult APCs for differentiation in the setting of obesity serves an important homeostatic role and thus phenotypic programming of the adult APC pool may influence responses to obesity. At sexual maturity, APCs isolated from iWAT of offspring of obese dams had higher adipogenic competency and lower proliferation rates *in vitro* (Fig. 5A – G), suggesting an enrichment in committed APCs. This high adipogenic/low proliferative phenotype was more striking in females, possibly due to earlier pubertal maturation in females (Fig. 5G). To determine if the higher intrinsic adipogenic potential translated to the *in vivo* setting, adipogenesis was pharmacologically stimulated in adult offspring by administration of rosiglitazone (rosi) for 8 weeks (Fig. 5H). In a recent study from our group revealing greater capacity for hyperplastic expansion of subcutaneous depots in female mice, we treated mice with rosi or vehicle-control for 8 weeks and showed that body fat mass gain in response to rosi predominated in females and was driven by expansion of subcutaneous depots [14]. Interestingly, females were protected against the exhaustion in APCs observed in males after stimulation of adipogenesis by rosi [14], substantiating findings by Graff *et al* [27]. These results suggest that the capacity for APC self-renewal is limited even in the absence of obesity, making rosi a good tool to determine the impact of the intrauterine environment on APC responses and self-renewal capacity. In the current study, male offspring of obese dams gained significantly more body fat when treated with rosi compared to offspring of lean dams (Fig. 5I). Rosi-induced depletion in CD34^+^/PDGFRα^+^ APCs was more severe in male offspring of obese dams, and this was observed exclusively in the iWAT depot (Fig. 5J). Since there were no differences in APC abundance prior to initiation of treatment, the difference in progenitor abundance between offspring groups at the end of the 8 week rosi treatment is likely a result of stimulation of adipogenesis by rosi (Fig. 5M). In contrast, APCs were enriched in both iWAT and vWAT of rosi-treated female offspring born to obese vs. lean dams, despite no difference in body fat accumulation (Fig. 5K, L). Overall, these results show that intrauterine exposure to maternal obesity programs an adult APC pool that is hyper-responsive to adipogenic stimuli, predisposing males with limited self-renewal capacity to a depletion in the APC pool.

**Figure 5.**
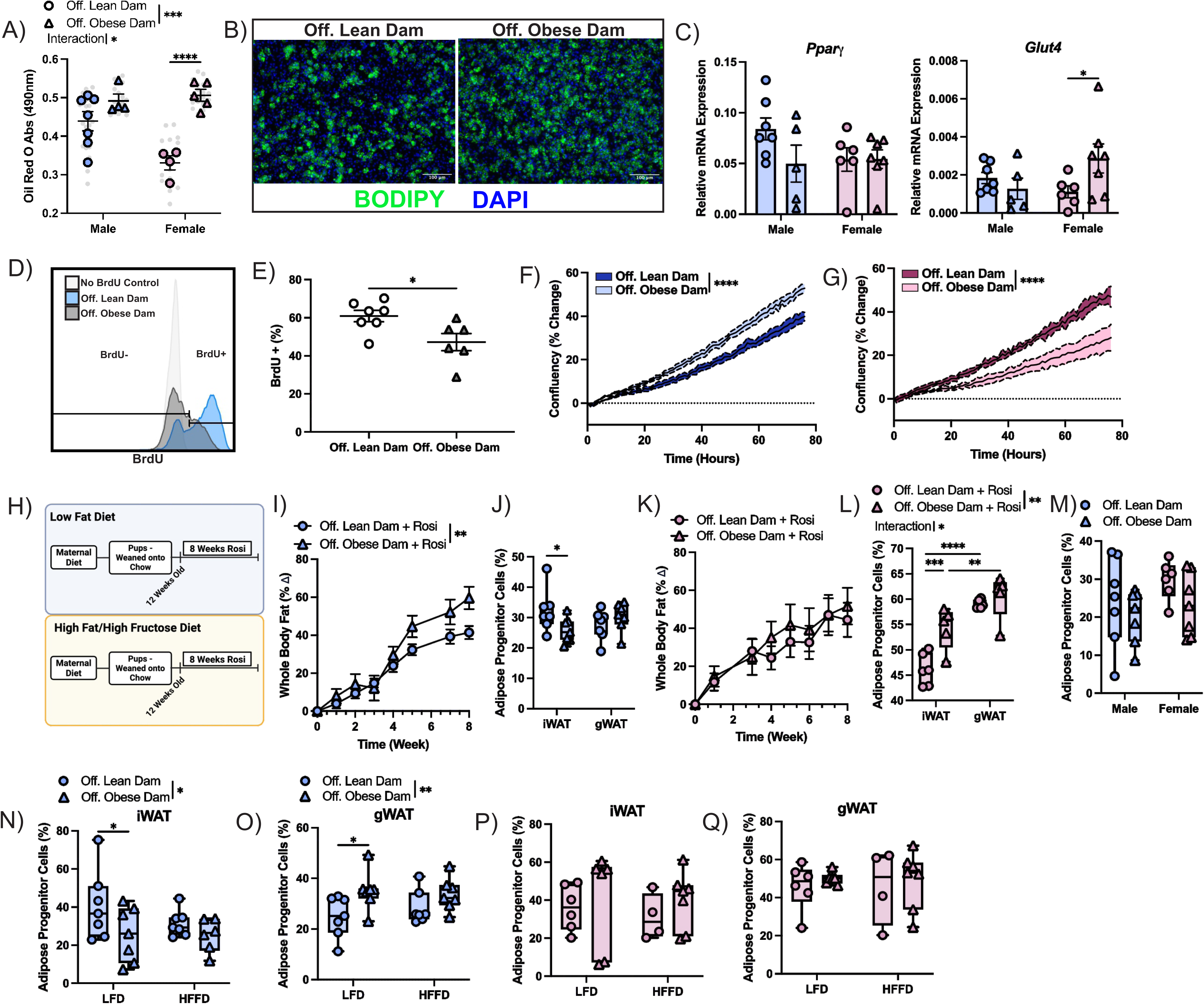
APCs Derived from Offspring of Obese Dams Lose Capacity for Self-Renewal in Adulthood. A) Quantification of neutral lipid staining by Oil Red O in APCs derived from sexually mature offspring of lean vs. obese dams after 7 days of *in vitro* differentiation. B) Representative images of *in vitro* differentiation with BODIPY staining. C) Expression of adipogenic genes *Pparγ* and *Glut4* after 7 days of *in vitro* differentiation. D) Gating paradigm for identification of BrdU^+^ APCs derived from adult offspring of lean vs. obese dams cultured *in vitro*. E) Flow cytometric quantification of BrdU^+^ APCs after overnight incubation with BrdU. F) Percent change in confluency of male and (G) female APCs derived from offspring of lean vs. obese dams, imaged on the Incucyte Zoom system for 72 hours. H) Experimental paradigm for administration of rosiglitazone. Offspring of lean or obese dams were administered rosi daily for 8 weeks beginning at 12 weeks of age. I) Percent change in whole-body fat of male offspring over 8 weeks treatment with rosi. J) APC abundance in iWAT and gWAT after 8 weeks treatment with rosi in male offspring of lean vs. obese dams. K) Percent change in whole-body fat of female offspring over 8 weeks treatment with rosi. L) APC abundance in iWAT and gWAT of female offspring after 8 weeks of treatment with rosi. M) iWAT APC abundance in male and female offspring prior to initiation of rosi. N) APC abundance in (N) iWAT and (O) gWAT of male offspring after 16 weeks of LFD or HFFD. APC abundance in (P) iWAT and (Q) gWAT of female offspring after 16 weeks of LFD or HFFD. *p<0.05, **p<0.01, ***p<0.001, ****p<0.0001.

Previously published work has shown an activation of adipogenesis in the setting of obesity in a depot and sex-dependent manner such that APC responses in males predominate in the visceral fat, whereas as APCs respond in both depots of females [14, 28]. Therefore, we also determined the effect of 16-weeks of LFD or HFFD feeding on progenitor abundance by flow cytometry. Male offspring of obese dams fed LFD had significantly fewer APCs than offspring of lean dams, an effect which was restricted to the iWAT depot (Fig. 5N). In the gWAT depot, progenitor abundance was increased in male offspring of obese relative to lean dams on LFD (Fig. 5O). In HFFD-fed males, there were no differences in APC abundance dependent on maternal diet in either depot (Fig. 5N, O). Female offspring displayed no differences in progenitor abundance as a result of either LFD or HFFD feeding in either depot (Fig. 5P, Q). These results provide further evidence for predisposition to APC exhaustion in the iWAT of males born to obese dams.

### Offspring of Obese Dams Have Exaggerated Susceptibility to Metabolic Consequences of Diet-Induced Obesity

We next sought to determine whether offspring of obese dams were more vulnerable to metabolic dysfunction. The frequency of large adipocytes (>100μm) in male iWAT was significantly higher in offspring of obese dams fed a HFFD (Fig. 6A – C). Comparatively, there were no differences in iWAT adipocyte size attributable to maternal or offspring diet in female offspring (Supp. Fig. 4A, B). Excessive obesity-related adipocyte hypertrophy in males may be due to the loss of available APCs for differentiation. In male gWAT, adipocyte size was primarily determined by diet (Fig. 6 – F). In female offspring gWAT, there were fewer small (<40μm) and more large (>100μm) adipocytes in offspring of obese dams fed a LFD and HFFD, respectively (Supp. Fig. 4C, D).

**Figure 6.**
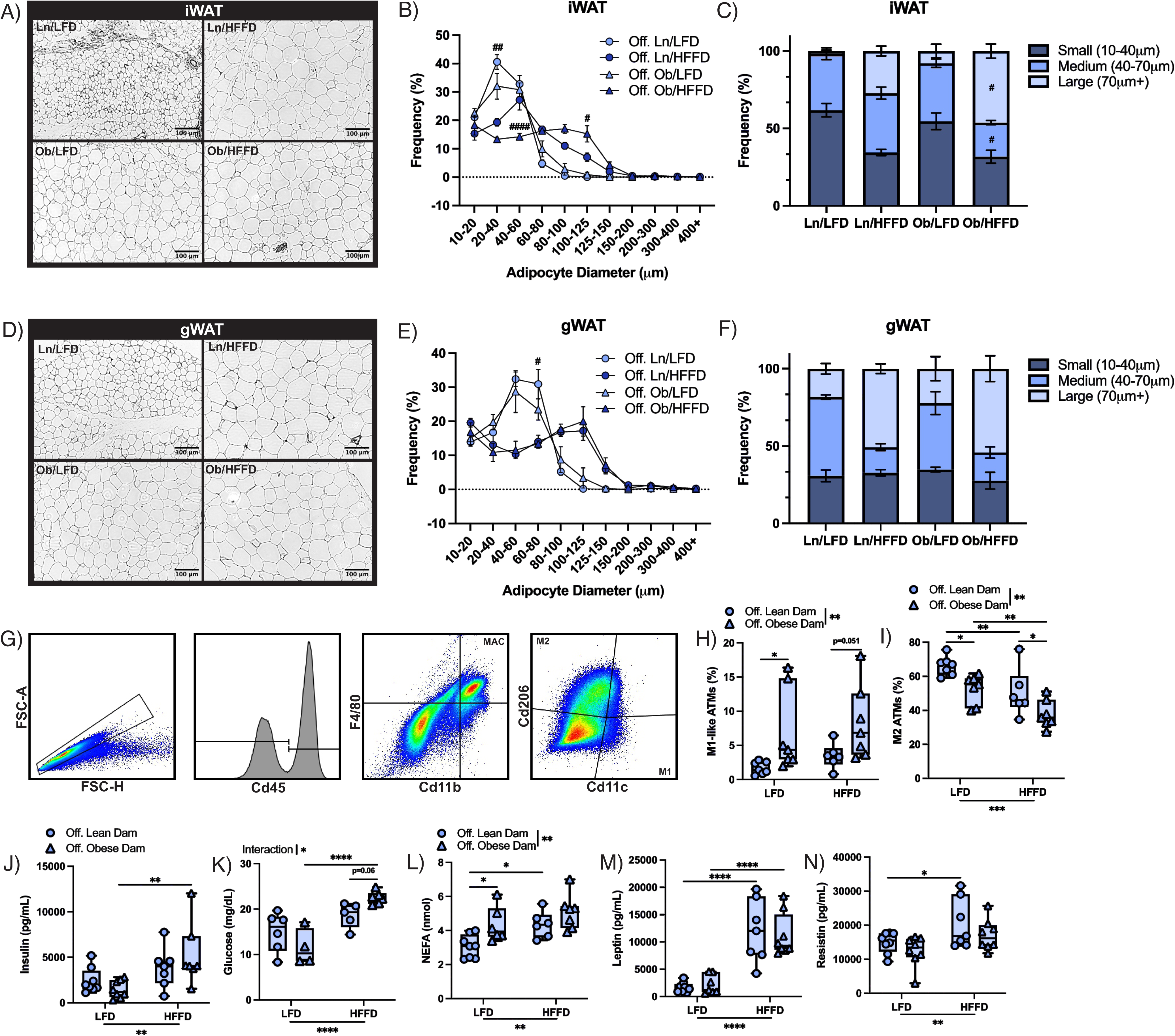
Offspring of Obese Dams are Vulnerable to Adipose Tissue and Systemic Metabolic Dysfunction. A) Representative images of H&E stained iWAT sections from male offspring after 16-weeks LFD or HFFD. B, C) Adipocyte size distribution in iWAT of male offspring after 16-weeks LFD or HFFD. D) Representative H&E-stained gWAT sections from male offspring after 16-weeks LFD or HFFD. E, F) Adipocyte size distribution in gWAT of male offspring after 16-weeks LFD or HFFD. G) Gating paradigm for quantification of M1 and M2 adipose tissue macrophages in gWAT of male offspring after 16-weeks LFD or HFFD. H) Quantification of M1 and (I) M2 macrophages in offspring gWAT after LFD or HFFD feeding. J) Serum insulin, (K) glucose, (L) NEFA, (M) leptin, and (N) resistin (M) in offspring after 16-weeks LFD or HFFD feeding. *p<0.05, **p<0.01, ***p<0.001, ****p<0.0001. #p<0.05 between offspring groups on same diet.

To evaluate WAT inflammation, a hallmark of WAT dysfunction, the abundance of M1-like and M2-like macrophages in the gWAT of male offspring was determined by flow cytometry (Fig. 6G). On both LFD and HFFD, offspring of obese dams had a greater proportion of M1-like macrophages (Fig. 6H). M2-like macrophages were decreased in offspring of obese relative to lean dams on either a LFD or HFFD, and HFFD decreased the proportion of M2-like macrophages in both groups relative to LFD (Fig. 6I). Features of systemic metabolic dysfunction, including serum insulin and fasting glucose were higher in HFFD-fed male offspring of obese vs. lean dams (Fig. 6J, K). Circulating non-esterified fatty acids (NEFA), a marker of WAT dysfunction, were higher in LFD-fed male offspring of obese dams vs. lean dams, with no further increase on HFFD (Fig. 6L). Leptin was increased as a result of HFFD in both groups, likely reflecting adiposity, but there were no differences in leptin levels within diet groups (Fig. 6M). Fasting glucose in females was unaffected by offspring group or postnatal diet (Supp. Fig. 4E). These data confirm that male offspring of obese dams have greater vulnerability to metabolic dysfunction when subjected to an obesogenic diet.

## DISCUSSION

The premise behind the developmental origins of health and disease (DOHaD) is that the fetus undergoes adaptations to a suboptimal intrauterine environment to ensure short-term survival at the cost of lifelong health. Much of this plasticity that affords the fetus the ability to adapt its developmental trajectory to fit the environment is lost in the postnatal period. Thus, these altered trajectories during sensitive windows of development lead to persistent changes in organ function. Animal studies have identified perturbations in myogenesis, nephrogenesis and cardiomyocyte maturation as key developmental mechanisms underlying later-life risk for cardiometabolic disease in fetuses growing in an oxygen or nutrient-deprived environment [29–33]. In addition to these adaptations in trajectories of cellular proliferation and differentiation that preserve limited resources for high priority organs, programming of neuroendocrine axes in a nutrient-scarce environment enhances efficiency of energy storing mechanisms, creating a mismatch with a postnatal environment of overnutrition and thereby vulnerability to obesity.

What remains unclear are the developmental mechanisms that give rise to later cardiometabolic risk in offspring growing in an intrauterine environment of nutrient overabundance and born at the other end of the birth weight spectrum. Given that high adiposity is a characteristic feature of macrosomic babies born to mothers who were obese or diabetic during pregnancy, it is likely that the primary developmental adaptation to an intrauterine environment of nutrient overabundance occurs in adipose tissue. However, direct impacts on adipose organogenesis have been largely decentralized in the DOHaD literature in favour of neuroendocrine explanations. Proposed neuroendocrine mechanisms attribute increased adiposity in the offspring to hyperphagia, occurring as a result of a perturbation in the maturation of appetite-regulating hypothalamic circuits, leading to leptin resistance [34]. However, the rodent models of maternal obesity used in these studies and the vast majority of studies produce growth restricted fetuses, apparently due to placental dysfunction. We undertook a pilot study to test various diet durations and determined that 8 weeks of pre-gestational exposure to the obesogenic diet resulted in fetal growth restriction, whereas growth restriction was not observed with the 4 weeks of diet used for the current study. In the growth restricted fetus, neuroendocrine programming of enhanced food intake is a mechanism contributing to subsequent early life catch-growth and later-life vulnerability to obesity. Reprogramming of satiety pathways and other adaptations that prioritize resource guarding in the setting of a nutrient-scarce intrauterine environment fit with the thrifty phenotype hypothesis that posits metabolic adaptations of the fetus prepare the fetus for its predicted extrauterine environment. We propose that the fetus exposed to excess glucose and lipids from obese or diabetic mothers adapts to its environment of nutrient overabundance by accelerating the development of subcutaneous adipose tissue, the body’s primary site of energy storage and lipid buffering organ. Indeed, fetal overgrowth leading to large-for-gestational age infants with excessive adiposity is by far the most common outcomes of pregnancies complicated by maternal obesity or diabetes, while growth restriction is relatively rare.

The sensitive window of adipose development spanning from late gestation into early postnatal life coincides with a period of APC fate determination [13] and APC proliferation [35] which is followed by a wave of adipogenesis and fat accumulation. Data from the current study demonstrate that by postnatal day 10, offspring of obese dams have markedly higher adiposity that is not accounted for by increased adipocyte size. Rather, scRNA-seq data revealed a relatively higher proportion of proliferative APCs and this transcriptomic data was substantiated by *in vitro* experiments showing a hyperproliferative phenotype. Thus, the fetus adapts to an *in utero* environment of nutrient over-abundance by enhancing energy storage capacity with an increase in adipocyte number achieved through extending the phase of APC proliferation during the critical window of adipose organogenesis that spans late gestation into the early postnatal phase. This APC hyperproliferation may be the result of maternal hyperglycemia, which elicits fetal hyperinsulinemia, a nutritional cue known to regulate adipose tissue development [36, 37]. A study by Yuan *et al.* determined that insulin signaling is required for proliferation of adipose progenitor cells at postnatal day 10 [38], the period during which developmental adipogenesis is declining and a greater proportion of resident APCs belong to the adult APC compartment [13]. Thus, it is likely that excessive fetal hyperinsulinemia is a trigger for APC hyperproliferation in pregnancies complicated by maternal obesity. Overall, our data reveal hyperproliferation of APCs and an endowment of adipose tissue with a greater number of adipocytes as the developmental origin of early-onset obesity in fetuses adapting to greater energy storage demands in pregnancies complicated by maternal obesity. Our findings align with a hallmark study from Spalding *et al.* demonstrating humans with higher adipocyte number in early life will have higher adiposity throughout the lifespan [10].

Our results show that the higher setpoint of adiposity in offspring exposed to maternal obesity *in utero* manifests in early postnatal life and prior to the onset of puberty. While there is very little known about adipogenesis in puberty, evidence from Holtrup *et al.* determined that a second peak of APC proliferation occurs, which corresponds to a second period of fat accumulation. However, it is unknown whether the pubertal surge in fat accumulation is attributable to differentiation of these proliferating APCs, or if adult APCs undergo proliferation to establish a reservoir of APCs for adulthood. We observed a prepubertal plateau in body weight in control offspring that was absent in offspring of obese dams. While offspring of obese dams continued to gain body mass in the pre-pubertal period, the pubertal surge in adiposity was absent. While a thorough investigation of pubertal adipose development is outside the scope of the current study, it is possible that the expected surge in adiposity was merely imperceptible given that offspring of obese dams had remarkably higher pre-pubertal adiposity when compared to controls. Further, there is some evidence that fat mass in and of itself acts as a trigger for puberty, and thus puberty and the corresponding fat surge may have occurred earlier. While a follow up study is required to fully determine the mechanisms underlying these pubertal differences in fat accumulation, these data nonetheless demonstrate that this important secondary period of adipose development is perturbed by exposure to maternal obesity during development.

The ‘Adipose Tissue Expandability Hypothesis’ posits that rather than absolute fat mass, the capacity for adipogenesis determines metabolic disease risk in the context of chronic caloric overload. Given that both developmental and adult progenitor cells are established during intrauterine life, each are subject to the programming influence of maternal obesity [13]. Thus, an intrauterine environment of nutrient excess may not only raise the setpoint of adiposity but also influence the plasticity of adipose tissue in later life. Our data demonstrate that following puberty, APCs derived from offspring of obese dams have increased adipogenic potential but decreased proliferative capacity. There is evidence from a number of organ systems demonstrating that self-replicative capacity of progenitor or stem cells has a lifetime limit. Thus, these data suggest that the pro-adipogenic phenotype of adult APCs may be a factor in the vulnerability to obesity in adulthood and lead to a premature decline in self-renewal capacity of the APC pool. Evidence from Tang *et al.* [27], our group [14], and others [26, 39] have suggested that the pool of adipose progenitor cells is finite and capable of being exhausted. Interestingly, Tang *et al.*, demonstrated that chronic stimulation of adipogenesis by rosiglitazone resulted in a reduction in APC availability without an increase in APC death [27]. Rosiglitazone belongs to the thiazolidinediones (TZDs), a family of anti-diabetic drugs and synthetic ligands and potent agonists of PPARγ, the major transcriptional regulator of adipogenesis. The insulin sensitizing actions of TZDs are in part attributable to activation of adipogenesis [40]. Thus, TZDs are a useful tool for studying the impact of stimulating adipogenesis. We hypothesized that if offspring of obese dams were vulnerable to APC exhaustion, administration of TZDs would unveil this predisposition. Following chronic administration of TZDs, offspring of obese dams had a significantly reduced abundance of APCs when compared to offspring of lean dams. This occurred concomitantly with an increase in body fat, suggesting that there was an adipogenic response to rosiglitazone which facilitated an increase in body fat, but resulted in reduced progenitor availability. We also observed a reduction in APC abundance in adult offspring of obese dams when on a low-fat diet. In both the rosiglitazone and diet-fed animals, progenitor exhaustion was restricted to the subcutaneous adipose tissue. This is unsurprising, given that in the mouse, the subcutaneous adipose tissue develops *in utero*, while visceral adipose tissue development is a postnatal event. This further confirms that the effects on progenitor abundance is a likely consequence of *in utero* programming, given the apparent lack of effect in the visceral adipose tissue. Recent studies have provided strong evidence demonstrating that a lack of adipogenesis is correlated with a reduction in insulin sensitivity [11] and that genetically manipulating PPARγ expression to increase adipogenesis in the context of diet-induced obesity abrogates the systemic metabolic consequences [12]. Accordingly, in our model, offspring of obese dams demonstrated increased susceptibility to the metabolic effects of diet-induced obesity, including increased macrophage infiltration and adipocyte size and systemic metabolic dysfunction including hyperinsulinemia and hyperglycemia. Taken together, these data suggest that the premature exhaustion of the APC pool predisposes offspring of obese dams to the metabolic effects of diet-induced obesity.

Prior to puberty, we observed few sex differences resulting from maternal programming. After puberty, female offspring of both groups demonstrate higher adiposity than males, as expected. Females were protected from both metabolic dysfunction and progenitor exhaustion in the setting of obesity, which we demonstrated in a recent study to be the result of sex hormone-regulated adipose progenitor cell responses. Many studies of developmental programming report sex differences, which are often attributed to differences in placental function and developmental timing, leading to sexually disparate outcomes to the effects of programming [41]. However, we propose that at least in the context of maternal obesity, males and females are equally susceptible to early-life effects, but female protection arises at puberty as a consequence of the cardiometabolic protective effects of estradiol.

Collectively, our data are the first to comprehensively investigate the effect of maternal obesity on APC function and abundance. We demonstrate that exposure to maternal obesity *in utero* programs early-onset obesity in the offspring through increased adipocyte number achieved through an extension in APC proliferation during the first wave of adipogenesis. Programming of the adult APC pool predisposes offspring to later impairment in self-renewal capacity and premature exhaustion of APCs. This may be a key mechanism linking maternal obesity to increased risk for metabolic syndrome in the offspring. Understanding the underlying mechanisms for obesity programming is critical as we move toward minimizing the transgenerational risk of obesity.

**Supplemental Figure 1. Pre-pregnancy Feeding with High-Fat/High-Fructose Diet Induces Obesity and Glucose Intolerance.** A) Pre-pregnancy whole-body fat percentage after 2 weeks of LFD or HFFD feeding. B) Whole-body fat mass in dams after pup weaning on PND21. C) Gestational weight gain in lean vs. obese dams. D) Blood glucose after 12-hour fast. E, F) Blood glucose in lean vs. obese dams 1-hour after refeeding. G) Litter size for lean vs. obese dams. *p<0.05, **p<0.01, ***p<0.001.

**Supplemental Figure 2. Unbiased APC Clustering.** A) Comparison of distribution of canonical cell type clusters between groups. B) Unbiased clustering of APCs. C) Heatmap of marker genes for curated APC clusters.

**Supplemental Figure 3. Group Comparison of Lineage Occupancy.** A) Proportion of cells in preadipocyte and anti-adipogenic lineage compared between groups. B) Violin plots of common markers for APC clusters. C) Group comparison of proliferative gene expression in APC cluster.

**Supplemental Figure 4. Female Offspring of Obese Dams are Protected from Progenitor Exhaustion and Metabolic Aberrations in Obesity.** A, B) Distribution of adipocyte size in iWAT of female offspring after 16-weeks LFD of HFFD feeding. C, D) Distribution of adipocyte size in gWAT of female offspring after 16-weeks LFD or HFFD feeding. E) Fasted blood glucose in female offspring at the end of the diet. *p<0.05, **p<0.01, ***p<0.001, ****p<0.0001.

