## Supplementary figures and images for "Hyper-proliferation of Adipose Progenitors During Developmental Adipogenesis Programs Higher Adipocyte Number and Early-onset Obesity in Offspring Born to Obese Dams"

### Supplementary Figure 1

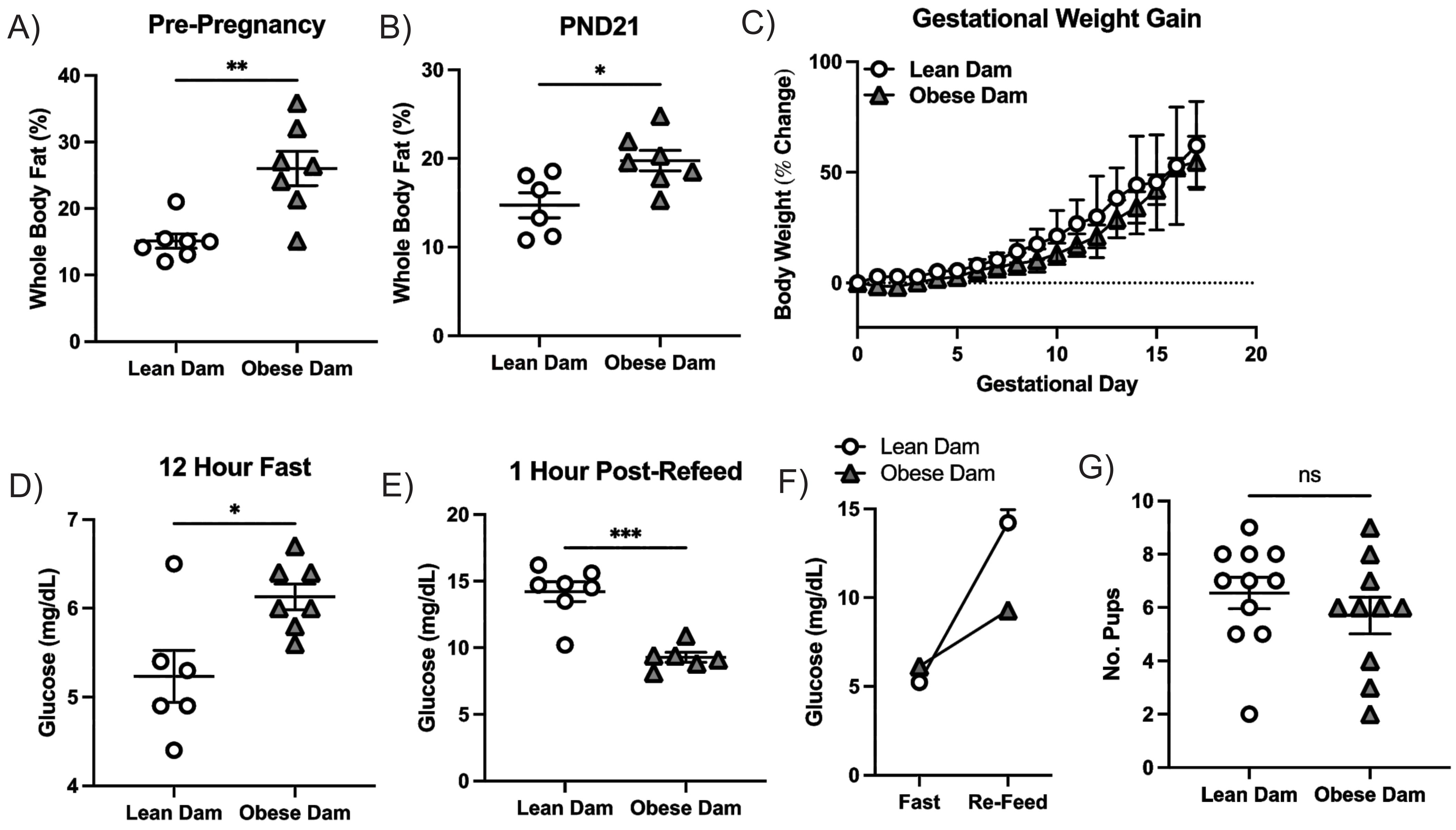

### Supplementary Figure 2

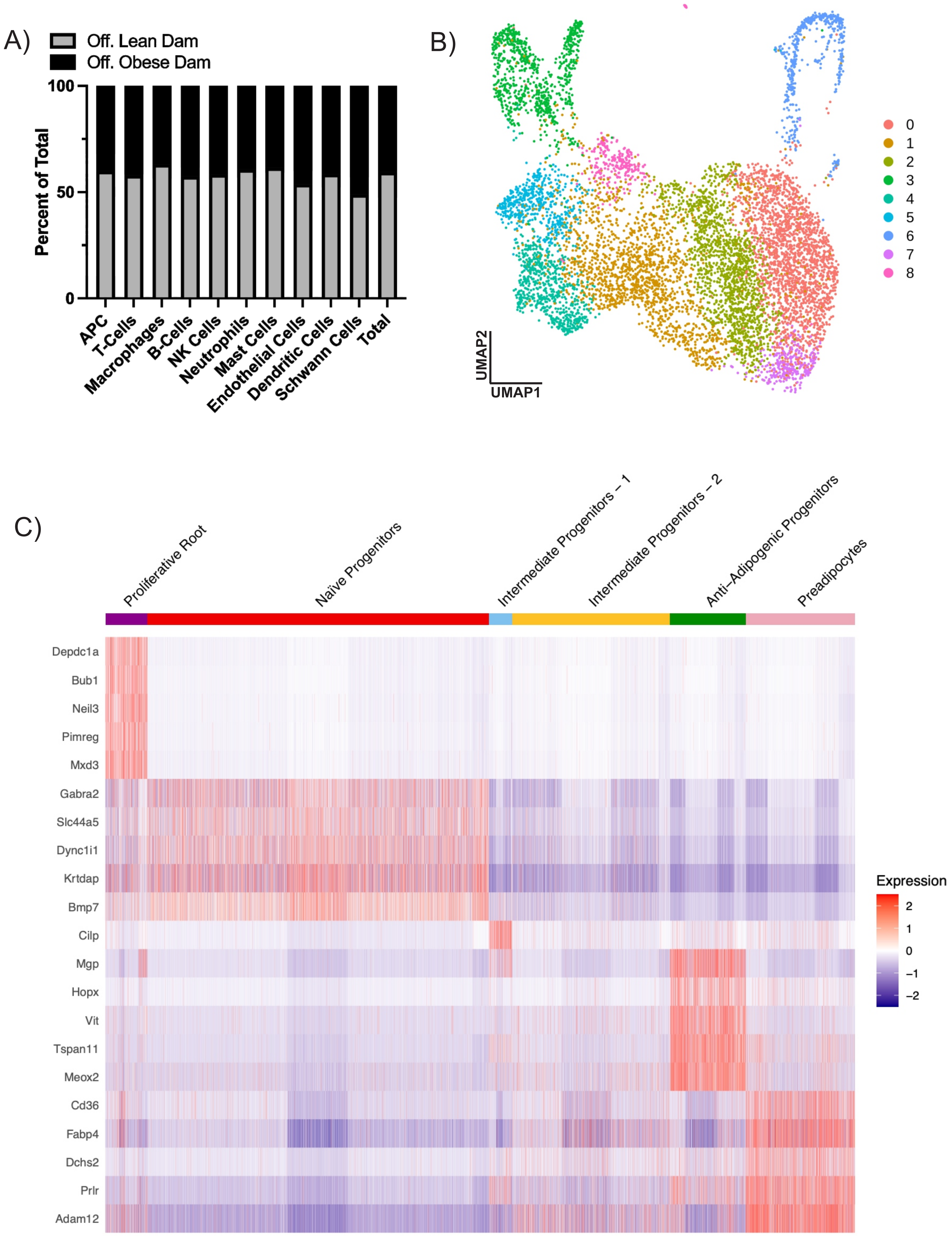

### Supplementary Figure 3

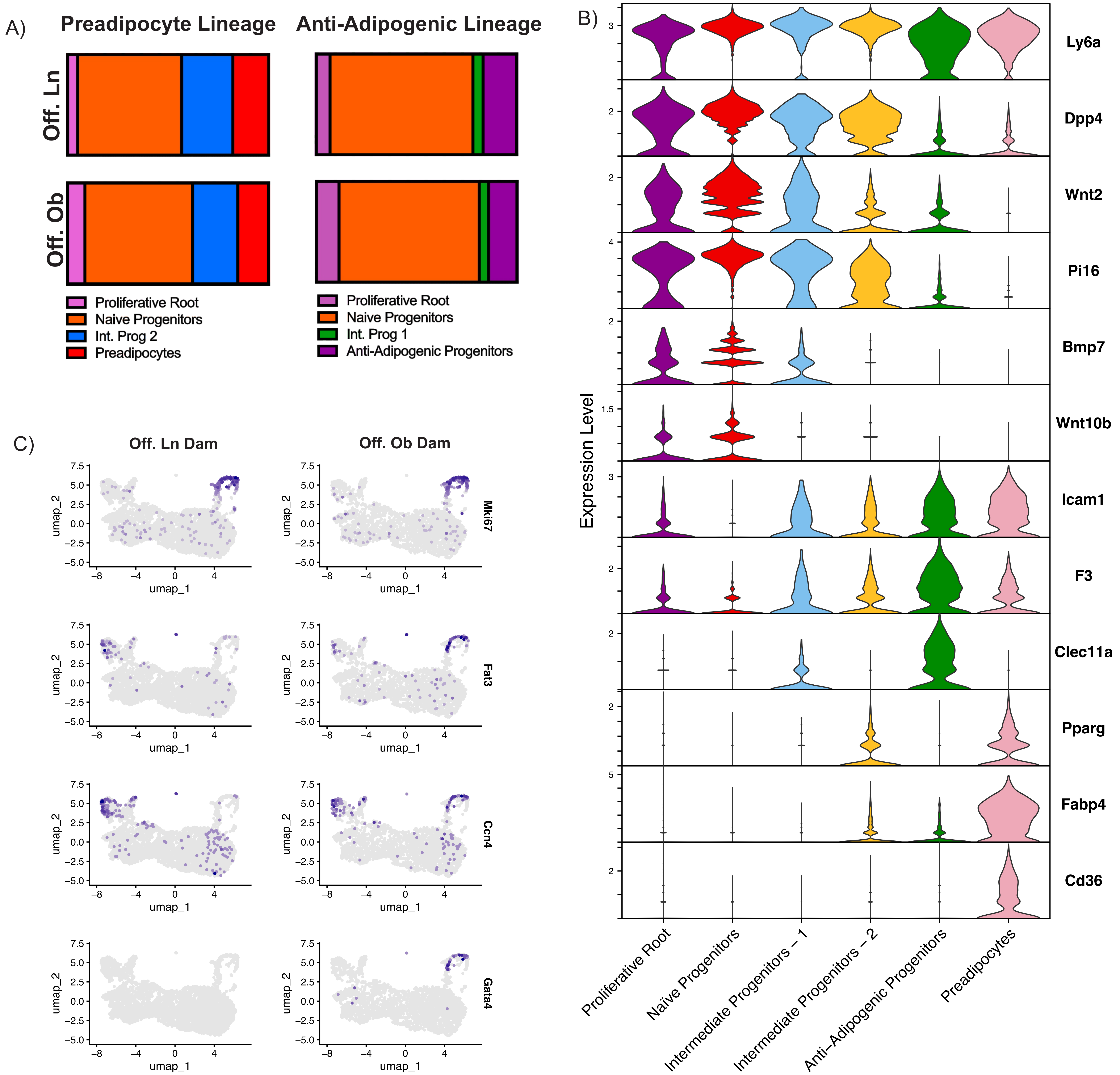

### Supplementary Figure 4

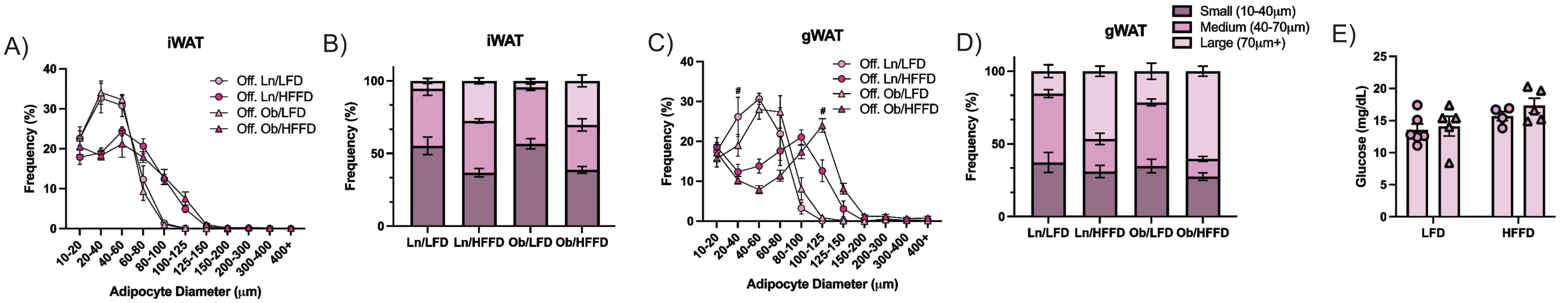
